# Detection of exotic biosecurity threat ribgrass mosaic virus and novel tobamoviruses through metatranscriptomic sequencing of animal gut content

**DOI:** 10.1101/2024.12.10.627875

**Authors:** Jackie Mahar, Jonathon C. O. Mifsud, Kate Van Brussel, Anna E. Lachenauer, Erin Harvey, Olivia M. H. Turnbull, Stefanie Bonat, Thomas M. Newsome, Annabelle Olsson, Antje Chiu-Werner, Menna E. Jones, Edward C. Holmes, Solomon Maina

## Abstract

Ribgrass mosaic virus (RMV) and related viruses of the genus *Tobamovirus* (*Virgaviridae*) are cruciferous plant pathogens that represent a threat to global horticultural systems. In Australia, they are considered exotic biosecurity threats, and an incursion of these viruses would require rapid and strict control efforts. However, current surveillance methods for these viruses are limited. We examined whether RMV and related tobamoviruses could be detected by deep sequencing of gut metatranscriptomes of vertebrate animals and ticks. Using this method, we discovered that RMV, as well as a novel relative of RMV, and two highly diverse novel tobamoviruses are present in Australia. RMV was detected in multiple sites in both the Australian Capital Territory (ACT) and Tasmania, two regions separated by approximately 700km of land and 200km of water. The novel relative of RMV was detected in the ACT and New South Wales (NSW), while the highly divergent novel tobamoviruses were each detected in a single state, NSW and Queensland (QLD). In addition, Tobacco mild green mosaic virus, which is already known to be present in Australia, was detected in QLD using this method. This work highlights the potential utility of metatranscriptomic sequencing of wild animal gut for the surveillance of biosecurity threats to native and agricultural plant species.

**Importance:** Plant viruses can have devastating impacts on global horticulture. Tobamoviruses (family *Virgaviridae*, genus *Tobamovirus*) are among the most damaging seed-borne viruses in horticultural crops, and Australia is free of many of the tobamoviruses that cause major crop losses in other countries. These viruses are extremely difficult to eradicate. Consequently, early detection of incursions is key to the control of these viruses in Australia, alongside rapid deployment of eradication and management plans. Current biosecurity surveillance methods in Australia rely on visual inspection, immunological assays, and molecular methods such as screening of imported seed lots. This study introduces a complementary approach that utilises unbiased metatranscriptomic sequencing of animal gut material to detect cryptic plant viruses circulating in nature. Using this approach, we detected five different tobamovirus circulating in Australia, including a virus thought to be exotic and three novel viruses. This unique approach highlights alternative options for surveillance/detection of exotic crop viruses.

## Introduction

Worldwide, virus disease outbreaks, epidemics and pandemics pose a threat to agricultural and horticultural systems (1). Viral diseases have been associated with impaired crop growth and vigour, leading to diminished or total loss of gross yields (2). This has been accelerated by agricultural globalization, climate change and other factors, such as pesticide resistance in virus vectors, that together are complicating disease management (3). Tobamoviruses (family *Virgaviridae*, genus *Tobamovirus*) are among the most damaging seed-borne viruses in horticultural crops (4). These viruses have a single stranded, positive-sense genome of about 6.3-6.5 kb that contain six open reading frames (ORFs): an RNA replicase read-through derivative (ORF1 and 2), an undefined protein (ORF3), movement protein (MP; ORF4), coat protein (CP; ORF5) (5), and occasionally an ORF6, that overlaps ORF4 and ORF5 (6). Tobamoviruses have an extensive host range and on this basis are divided into three subgroups: subgroup 1 infect solanaceous species, subgroup 2 infect Cactaceae, Cucurbitaceae, Fabaceae, Malvaceae and Passifloraceae species, and subgroup 3 infect a range of hosts, including Cruciferae/Brassicaceae and *Plantago* (Plantaginaceae) species (7, 8). Tobamoviruses have been associated with substantial economic losses linked to reduced crop yield as well as fruit damage which triggers a significant reduction in marketability (9, 10). For example, tomato brown rugose fruit virus (ToBRFV), is a damaging new tobamovirus that infects tomatoes and peppers that was first discovered in Jordan (11) and Israel (12), later spreading to many countries worldwide (13). Similarly, cucumber green mottle mosaic virus (CGMMV) is a recently introduced tobamovirus in Australia (14) that causes serious disease and damage in the *Cucurbitaceae*, such that it is one of the most economically important cucurbit viruses (15).

Agriculture of non-native crops in Australia commenced after colonization by Europeans in 1788 and in some regions, as recently as 60 years ago (16). Such a recent timescale, as well as the isolation of this large island nation, means that many viruses and virulent virus strains that co-evolved with these plants and cause damaging crop diseases in other parts of the world are absent or have only recently arrived in Australia. Human activities within the global trade in plants and plant products, mostly involving unknowingly infected seed or vegetative propagules, may spread viruses from other countries to Australia (3, 17, 18). Evidence that this may have already occurred in the Australian horticultural industry include recent detections of CGMMV and pepper vein yellows virus (14, 19). Alternatively, viruses can spill over from indigenous wild plant populations to managed cultivated populations (20–22). As agriculture and horticulture continues to expand in Australia, so too does the overlap of wild plant and crop habitat, increasing chances of virus spillover (23). As such, understanding the range of viruses that exist on wild vegetation and their interactions with local animal fauna is an important complementary route for managing crop viral diseases. Indeed, plant viruses may colonize new environments through grazing animal teeth, ingestion of plants, and subsequent passing of the viruses through the gut (24). However, very few studies have comprehensively explored persistent viruses in wild plants (23, 25).

Recently, we inadvertently detected ribgrass mosaic virus (RMV) in Australia in wild animal gut content while exploring animal viromes using metatranscriptomic (i.e. total RNA) sequencing (26). RMV is considered a biosecurity risk and was previously considered to be exotic to Australia, but is endemic in neighbouring New Zealand (27). Its natural hosts include members of the Cruciferaceae and Plantaginaceae, and it is known to have a broad host range compared to other tobamoviruses (28). RMV belongs to *Tobamovirus* subgroup 3, and is most closely related to youcai mosaic virus (YoMV), turnip vein clearing virus (TVCV) and wasabi mottle virus (WMoV) (29), all of which are exotic plant pathogens and Australian biosecurity risks (https://www.agriculture.gov.au/biosecurity-trade/pests-diseases-weeds/plant/national-priority-plant-pests-2019, accessed 21/10/24).

Tobamoviruses are highly contagious, easily spread through mechanical transmission between plants and fomites (30), and in some cases are seed transmitted. Additionally, they can remain viable in soil for many months (even up to a year) and are very difficult to eradicate once present (31). As there are no available treatments, plant destruction and soil decontamination are the only means of control (32). Consequently, early detection and prevention of incursion is key to the control of these viruses in Australia, alongside rapid deployment of eradication and management plans in the event of incursions. Current biosecurity surveillance methods in Australia rely on visual inspection, immunological assays, and molecular methods such as screening of imported seed lots (32). Efforts to include targeted plant viral genomics are in development (33).

Herein, we explored the use of an alternative surveillance strategy, based on the metatranscriptomic sequencing of animal gut content, to unravel unknown circulating biosecurity threats within wild or cultivated vegetation. This approach was used (i) to determine the presence of biosecurity threat tobamoviruses in Australia, at the same time assessing the utility of this method, and (ii) to reveal the phylogenetic relationships among the tobamoviruses detected and their implications for the biosecurity of the Australian horticultural industry.

## Materials and Methods

### Metatranscriptomic sequence data

Metatranscriptomic data sets previously generated from studies of Australian animal viromes were opportunistically screened for the presence of tobamoviruses. These data sets were generated by sequencing total RNA extracted from whole ticks, and mammalian gut content in the form of faecal samples, caecal content samples, or anal/rectal swabs (Table 1). Total RNA was extracted using the Maxwell 16 LEV simply RNA tissue kit in combination with a Maxwell nucleic acid extraction robot (Promega, WI, USA) for rabbit caecal content; the Qiagen RNeasy Plus Universal Mini Kit (Hilden, Germany) for deer and kangaroo carcass swabs; and the Qiagen RNeasy Plus Mini Kit (sometimes in conjunction with Qiagen QiaShredder columns) for dog swabs, ticks and marsupial carnivore faecal samples. The rabbit RNA libraries were prepared using the TruSeq Total RNA library preparation protocol (Illumina, CA, USA), with rRNA removal using Illumina Ribo-Zero gold epidemiology rRNA removal kit. The Illumina Stranded Total RNA Prep with Ribo-Zero Plus kit was used for all the other libraries. Paired-end sequencing was conducted on an Illumina HiSeq 2500 (200 cycles) for the rabbit libraries, and Illumina NovaSeq 6000 S4 lane (300 cycles) was used for all the other libraries) (Illumina, CA, USA). All library preparation and sequencing was conducted at the Australian Genomic Research Facility (AGRF). The raw reads obtained were trimmed using Trimmomatic v0.38 (34) and *de novo* assembled using MEGAHIT v1.2.9 (35) or Trinity v2.5.1 (36). Further details on data generation and ethics for data sets collected from rabbits can be found in Mahar et al. 2020 (26). Manuscripts are in preparation for the remaining studies.

**Table 1.** Metadata of metatranscriptome libraries containing tobamoviruses.

| Collection location | Library name | Animal species (common name) | Sample type | No. samples in pool | D.O.C | Tobamovirus(es) detected |
| --- | --- | --- | --- | --- | --- | --- |
| Crackenback/<br>Moonbah, NSW | 12-Kang2B | *Macropus giganteus*<sup>b</sup><br>(Eastern grey kangaroo) | Mouth and anal swab | 4 | Feb 2021 | SG3-like tobamovirus |
| Crackenback/<br>Moonbah, NSW | 13-Deer2A | *Dama dama*<sup>b</sup> (Fallow deer) | Mouth and anal swab | 4 | Dec 2020 and Jan 2021 | SG3-like tobamovirus |
| Crackenback/<br>Moonbah, NSW | 15-Deer2B | *Dama dama*<sup>b</sup> (Fallow deer) | Mouth and anal swab | 4 | Jan 2021 | Bambi tobamovirus |
| Crackenback/<br>Moonbah, NSW | 16-Deer2B | *Dama dama*<sup>b</sup> (Fallow deer) | Mouth and anal swab | 5 | Feb and Mar 2021 | SG3-like tobamovirus, Bambi tobamovirus |
| Hope Vale, QLD | HPV3-R | *Canis lupus familiaris*<br>(Domestic dog) | Rectal swab | 2 | Dec 2021 | Bluey tobamovirus |
| Hope Vale, QLD | HPV4-R | *Canis lupus familiaris*<br>(Domestic dog) | Rectal swab | 3 | Dec 2021 | Bluey tobamovirus |
| Hope Vale, QLD | HPV6-R | *Canis lupus familiaris*<br>(Domestic dog) | Rectal swab | 1 | Dec 2021 | Bluey tobamovirus, TMGMV |
| Hope Vale, QLD | HPV15-2R | *Canis lupus familiaris*<br>(Domestic dog) | Rectal swab | 2 | Feb 2022 | Bluey tobamovirus |
| Hope Vale, QLD | HPV16-2R | *Canis lupus familiaris*<br>(Domestic dog) | Rectal swab | 3 | Feb 2022 | Bluey tobamovirus |
| Hope Vale, QLD | HPV35-2E | *Rhipicephalus sanguineus* (tick),<br>attached to domestic dog | Whole animal | 1 | Feb 2022 | Bluey tobamovirus |
| Ben Lomond, TAS | P04 | *Dasyurus maculatus*<br>(Spotted-tailed quoll) | Faecal | 2 | Apr-Aug 2021 | RMV |
| Northern Slopes, TAS | P19 | *Felis catus* (feral cat) | Faecal | 1 | Apr-Aug 2021 | RMV |
| Northern Slopes,<br>TAS | P21 | *Dasyurus maculatus*<br>(Spotted-tailed quoll) | Faecal | 1 | Apr-Aug<br>2021 | RMV |
| Northern Slopes,<br>TAS | P22 | *Sarcophilus harrisii*<br>(Tasmanian devil) | Faecal | 5 | Apr-Aug<br>2021 | RMV |
| Crace, ACT | GUN-CC<sup>a</sup> | *Oryctolagus cuniculus*<sup>b</sup><br>(European rabbit) | Caecal<br>content | 20 | Dec 2016-Jan 2017 | RMV, SG3-like<br>tobamovirus |
| Booth, ACT | Gudg-CC<sup>a</sup> | *Oryctolagus cuniculus*<sup>b</sup><br>(European rabbit) | Caecal<br>content | 18 | Feb 2017 | RMV |

a - published in Mahar et al., 2020
b – samples taken from carcasses, not live animals.

### Ethics

For deer and kangaroo carcasses, scientific licenses and collection permits were obtained to relocate carcasses (SL102334) and research was approved by the University of Sydney Animal Ethics Committee, project number: 2019/1640. Dog and tick sampling was approved by the Taronga Conservation Society Australia’s Animal Ethics Committee, approval number 4c/10/21. Ethics for Tasmanian carnivores was obtained from the University of Tasmania under Animal Ethic Committee permit number A0018012. Ethics for rabbit sampling is as described in Mahar et al., 2020 (26).

### Tobamovirus detection

The assembled contigs were annotated using DIAMOND BLASTx (37), and blastn (38) against NCBI’s non-redundant protein and nucleotide databases, respectively. Contigs with top hits to tobamovirus were selected for further analysis. Open reading frames were identified using the Find ORFs tool within Geneious Prime ® 2022.2.2 using the standard genetic code and UAG read-through. ORFs were adjusted and annotated based annotations copied from related tobamoviruses using the Annotate from Database tool available within Geneious Prime. ORF gene assignments were confirmed by identification of conserved domains using RSP-TBLASTN v2.6.0 (39) and the NCBI CDD database.

### Phylogenetic classification of tobamoviruses

Contigs that had a top blast hit to a tobamovirus were confirmed through sequence alignment and phylogenetic analysis with other published tobamovirus sequences. Accordingly, the tobamovirus NCBI nucleotide RefSeqs, the top blast hits to the tobamoviruses detected here, as well as all published nucleotide sequences of RMV, TVCV, WMoV, YoMV, and TMGMV were downloaded from the NCBI nucleotide database (https://www.ncbi.nlm.nih.gov/) and aligned with the tobamovirus nucleotide sequences detected in this study using MAFFT v7.490 (40). Ambiguously aligned regions were removed using trimAl v1.4.1 (41). Nucleotide substitution model selection was performed (42) and maximum likelihood phylogenies inferred using IQ-TREE v2.1.3 (43). Branch supports were estimated using bootstrapping (1,000 replicates). Phylogenetic trees were inferred for both the complete genome (n=131, 8,341 nt), as well as the coat gene, the most commonly sequenced gene among tobamoviruses (n=167, 528 nt). A distance matrix for subgroup 3 tobamoviruses was inferred by aligning all available subgroup 3 nucleotide sequences from NCBI with the new sequences from this study, using MAFFT. Identical sequences were removed and genetic distances were calculated (as percentage nucleotide identity) and visualized in Geneious Prime ® 2022.2.2.

### Recombination detection

All new tobamovirus species were screened for the presence of recombination using the recombination detection program v4.96 (RDP4) (44), using the default parameters. A full genome alignment including the new tobamoviruses detected here, NCBI RefSeqs for subgroup 1 and 3 viruses, and the top blast hits for each tobamovirus contig was used as input (n=75 sequences). The highest acceptable p-value was set to 0.05 with Bonferroni correction, and recombination events were only considered if detected by at least three different methods.

### Meta-transcriptomic identification of potential plant host species

CCMetagen (45) was used to classify the non-viral transcripts in the libraries where tobamoviruses were detected, using the default method of abundance calculation. Classification was based on screening against the curated indexed database compiled by the CCMetagen creators, which contains the NCBI nucleotide collection, with the exception of most artificial and environmental sequences that lack taxids.

### PCR confirmation

The presence of the novel or exotic tobamoviruses detected within each metatranscriptomic library were confirmed by amplifying a small fragment of the relevant virus (RMV, 565 bp; SG3-like tobamovirus, 571 bp; bluey tobamovirus, 226 bp; bambi tobamovirus, 540 bp). For RMV, the SuperScript™ One-Step RT-PCR System with Platinum™ Taq DNA polymerase (Invitrogen, MA, USA) was used according to the manufacturer’s instructions. RMV isolate PV-051 infected lyophilized plant material imported from DSMZ Germany was used as a positive control. Optimum cycling conditions were as follows: 50°C for 30 min for reverse transcription, 95°C for 15 min followed by 35 cycles at 95°C for 30 s, 55°C for 40 s, and 72°C for 45 s, with a final extension at 72°C for 10 min. For all other viruses, the RT-PCRs were conducted in a different lab using the SuperScript^TM^ IV One-Step RT-PCR system (Invitrogen) according to the manufacturer’s protocol. The cycling conditions were as follows: 50°C for 10 mins and 98°C for 2 mins, followed by 35 cycles of 98°C for 10 s, 55°C for 10 s and 72°C for 30 s, with a final extension of 72°C for 5 mins. The amplified specific PCR products generated were confirmed by gel electrophoresis followed by SYBR safe staining. At least one set of PCR products for each virus were purified and confirmed by Sanger sequencing at the Australian Genome Research Facility (AGRF) at Westmead, NSW.

### Amplification and Sanger sequencing confirmation of SG3-like tobamovirus virus genome

To confirm the consensus genome sequence of SG3-like tobamovirus virus, RT-PCR and Sanger sequencing was performed on sample 12.9 from library 16-Deer2B. The tobamovirus sequence was amplified in four segments (∼1040 bp to ∼2100 bp) using the SuperScript^TM^ IV One-Step RT-PCR system (Invitrogen) following the manufacturer’s protocol and using four primer sets (Table S1). The cycling conditions were as follows: 50°C for 10 mins and 98°C for 2 mins, 35 cycles of 98°C for 10 s, 55°C for 10 s and 72°C for 1 min, and a final extension of 72°C for 5 mins. DNA amplicons were purified using the GenElute PCR clean-up kit (Sigma-aldrich, MO, USA) and sent for single direction sanger sequencing at AGRF at Westmead, NSW, Australia using the amplification primers plus additional sequencing primers (Table S1). The resulting chromatograms were trimmed and mapped to the original metatranscriptomic-assembled genome sequence using Geneious Prime version 2022.1.1, and a consensus generated.

## Data Availability

The consensus sequences for tobamoviruses assembled in this study have been submitted to NCBI/GenBank and assigned accession numbers XXXXX-XXXXX.

## Results

### Detection and classification of tobamoviruses in animal gut metatranscriptomes

Viral contigs related to tobamovirus species were detected by Blast analyses of the gut metatranscriptomes of Australian wild animals collected from four states/territories in eastern Australia; NSW, TAS, ACT and QLD. Phylogenetic classification based on complete genome sequences (Figure 1A) revealed that five different tobamovirus species were present in the animal metatranscriptomes sampled here: Tobacco mild green mosaic virus (TMGMV), two subgroup 3 viruses, and two distinct novel tobamoviruses. Tobamovirus contigs varied in length, but the complete coding region of each of the five tobamoviruses detected here was assembled from at least one library.

**Figure 1.**
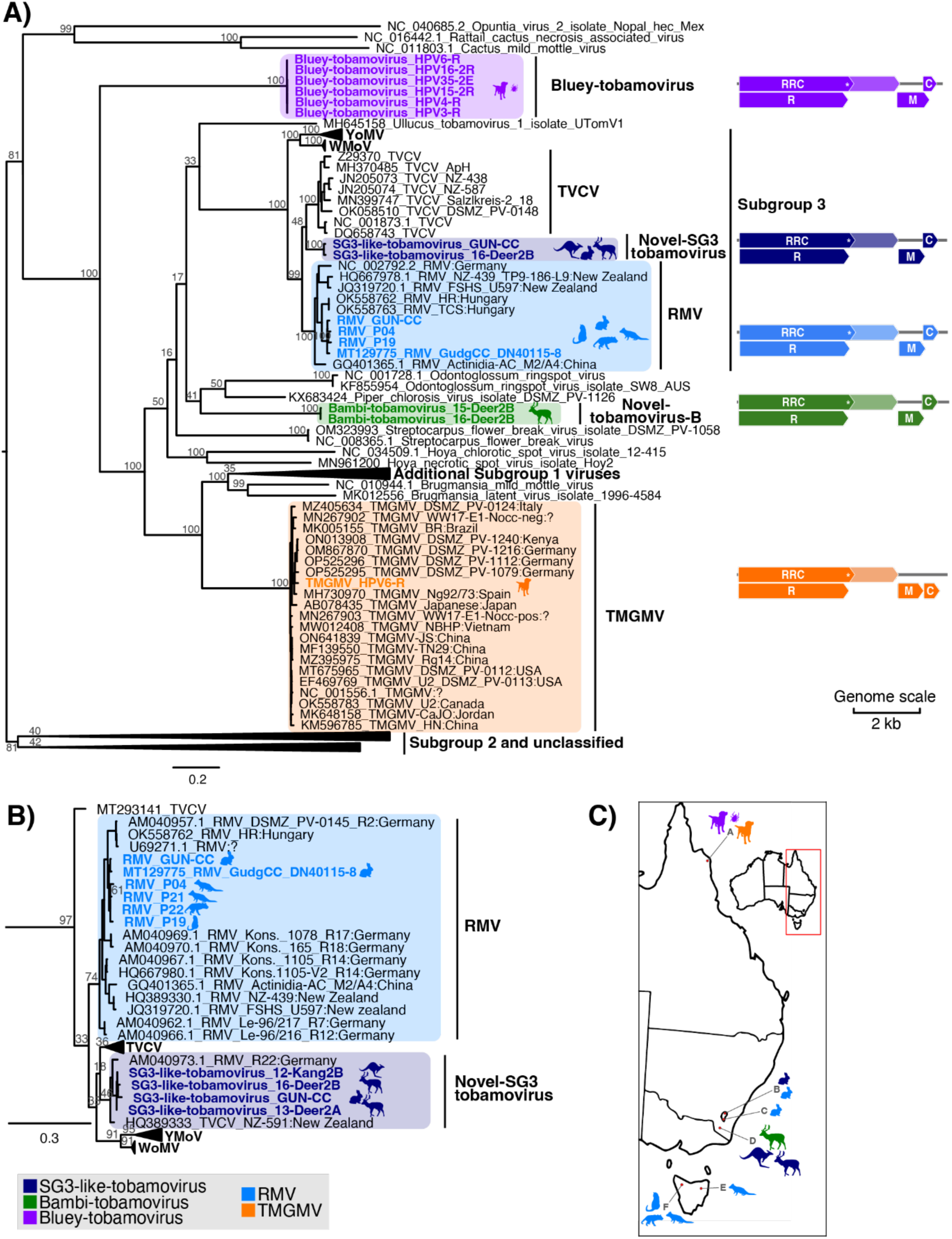
Diversity and geographic distribution of tobamoviruses found in animal meta-transcriptomes in Australia. **(A)** ML phylogenetic tree of the genome sequences of the tobamoviruses detected in Australian animal metatranscriptomes together with published tobamovirus genomes. **(B)** ML phylogenetic tree of the coat gene sequences of tobamoviruses detected in Australian animal metatranscriptomes together with published coat sequences, showing only the subgroup 3 clade (CP tree is in shown in Supplementary Figure 1). In the ML trees, sequences obtained from animal metatranscriptomes in Australia are indicated in bold and coloured by viral species, and the clades that they cluster within are highlighted in the same colour: purple=bluey tobamovirus; dark blue=SG3-like tobamovirus; blue=RMV; green=bambi tobamovirus; orange=TMGMV. Animal silhouettes beside the clades indicate the animal metatranscriptomes from which these viruses were obtained and are coloured by viral species. The GenBank accession number for published sequences is indicated at the start of the taxon name. Numbers at the nodes indicate the percentage support from 1,000 bootstrap replicates and the trees are midpoint rooted. The location (country) of collection is indicated in the taxa name in relevant clades. Genome schematics for viruses detected in this study are shown to the right of the relevant viral species, with coloured arrow-tipped boxes representing forward open reading frames for the following gene products: RNA replicase read-through component (RRC), RNA replicase (R), movement protein (M) and capsid protein (C). The read-through stop codon in the RRC is indicated by an asterisk and a break in the box, while the RNA-dependent RNA polymerase region translated via the read-through mechanism is indicated by lighter shading. **(C)** Map of eastern Australia indicating where tobamoviruses were detected. Animal silhouettes indicate the animal metatranscriptomes where these viruses were obtained and are coloured by viral species. Locations on the map are indicated by a letter and are as follows: A=Hope Vale, QLD; B=Crace, ACT; C=Booth, ACT; D=Crackenback/Moonbah, NSW; E=Ben Lomond, TAS; F=Northern Slopes, TAS.

Classification of the subgroup 3 viruses was hindered by the fact that many published virus sequences (particularly CP sequences) in the subgroup 3 clade have been misclassified. Despite this, it was possible to identify defined clades that represent each viral species *Tobamovirus plantagonis* (RMV), *Tobamovirus rapae* (TVCV), *Tobamovirus wasabi* (WMoV), and *Tobamovirus youcai* (YoMV) (Figure 1), each with 100% bootstrap support in the genome sequences tree. A distance matrix based on complete genome nucleotide sequences also supported that the species should be defined by these clades, in accordance with the ICTV species demarcation criteria for these viruses (https://ictv.global/report/chapter/virgaviridae/virgaviridae/tobamovirus), with sequences in the same clade sharing >90% similarity and those in separate clades sharing less than 90% nucleotide similarity (with a single outlier in YoMV) (Table 2).

**Table 2.** Nucleotide identity within and between tobamovirus subgroup 3 species.

|  | YoMV | WMoV | TVCV | RMV | SG3-like tobamovirus |
| --- | --- | --- | --- | --- | --- |
| **Maximum % identity (identical sequences have been removed)** |  |  |  |  |  |
| **YoMV** | 99.9<sup>a</sup> | 85.0 | 82.4 | 82.5 | 81.7 |
| **WMoV** | 85.0 | 99.7 | 82.7 | 83.3 | 82.6 |
| **TVCV** | 82.4 | 82.7 | 98.4 | 89.0 | 88.0 |
| **RMV** | 82.5 | 83.3 | 89.0 | 99.9 | 88.2 |
| **SG3-like tobamovirus** | 81.7 | 82.6 | 88.0 | 88.2 | 98.8 |
| **Minimum % identity** |  |  |  |  |  |
| **YoMV** | 89.1<sup>b</sup> | 83.1 | 80.6 | 79.5 | 80.5 |
| **WMoV** | 83.1 | 98.4 | 81.9 | 80.3 | 82.2 |
| **TVCV** | 80.6 | 81.9 | 92.6 | 85.4 | 87.2 |
| **RMV** | 79.5 | 80.3 | 85.4 | 90.3 | 86.0 |
| **SG3-like tobamovirus** | 80.5 | 82.2 | 87.2 | 86.0 | 98.8 |
| **Mean % identity** |  |  |  |  |  |
| **YoMV** | 94.2 | 84.5 | 81.6 | 81.0 | 81.3 |
| **WMoV** | 84.5 | 99.1 | 82.3 | 81.8 | 82.5 |
| **TVCV** | 81.6 | 82.3 | 94.0 | 87.0 | 87.6 |
| **RMV** | 81.0 | 81.8 | 87.0 | 93.6 | 86.7 |
| **SG3-like tobamovirus** | 81.3 | 82.5 | 87.6 | 86.7 | 98.8 |

a - Cells with >90% nucleotide identity are shaded grey
b - Minimum identity within YoMV sequences drops below 90% due to one sequence only

In clearly defining the subgroup 3 species clades, it was possible to determine that the two distinct SG3 viruses detected here belonged to RMV and a distinct novel SG3 group, tentatively named SG3-like tobamovirus (Figure 1A and B). The SG3-like tobamovirus clade contained sequences previously published as RMV or TVCV, including the complete genome of a virus detected by Mahar et al., 2020 (26) in rabbits (accession MT129780) and CP sequences from New Zealand (HQ389333) and Germany (AM040973) (Figure 1B). While phylogenetic analyses grouped these sequences with RMV and TVCV with strong bootstrap support, these viruses are distinct from both RMV and TVCV (Figure 1). Support for the SG3-like tobamovirus clade of viruses is not robust in the CP tree (bootstrap support of 46%; Figure 1B) due to limited informative sites, but the complete genome phylogeny strongly supports the existence of this small clade (Figure 1A). Furthermore, according to the *Tobamovirus* genus demarcation criteria (<90% nucleotide similarity between species) the SG3-like tobamovirus clade is diverse enough to constitute a new species, with all sequences in the SG3-like tobamovirus clade sharing no more than 88.2% identity with RMV and other viruses (Table 2), although this will need to be confirmed with additional data. It should be noted that because this virus was novel but closely related to RMV and TVCV, the metatranscriptome assembled genome sequence was confirmed by amplification and Sanger sequencing. The entire genome sequence was confirmed with the exception of a 395 nt region within ORF1 which could not be completed by Sanger sequencing due to sample insufficiency.

The other two novel viruses detected in this study, tentatively named Bluey Tobamovirus and Bambi Tobamovirus, were highly divergent from each other and all previously classified Tobamoviruses (Figure 1A). Bambi tobamovirus was most genetically similar to Piper chlorosis virus (PChV) isolate DSMZ PV-1126 (accession KX683424) based on nucleotide sequence identity but shared only 60.3% nucleotide similarity across the genome and these sequences did not cluster together (Figure 1). Similarly, bluey tobamovirus had the highest nucleotide identity to TVCV (accession OK058510), but shared only 57.5% nucleotide identity with this virus and did not form a monophyletic group with TVCV (Figure 1). These two highly novel viruses did not group into a known subgroup but were most closely related to subgroup 3 viruses. Finally, no viruses discovered here were determined to be recombinants.

### Distribution and diversity of detected tobamoviruses

TMGMV was detected in domestic dog rectal swabs from one location in QLD, while RMV was found in rabbit gut in two locations in the ACT, and in marsupial carnivore faeces in two locations in Tasmania (Figure 1C). Genetic diversity within the detected RMV viruses ranged from 97.4 – 99.3% nucleotide identity. SG3-like tobamovirus was detected in both the ACT in rabbit gut and in deer and kangaroo carcass swabs in NSW, sharing 98.1 – 99.9% nucleotide identity between them. Bluey tobamovirus was detected in a single location in QLD in five libraries sequenced from domestic dog rectal swabs as well as a library from a tick that was attached to a domestic dog. The bluey tobamovirus viruses discovered here shared 98.1 – 100% nucleotide identity, including the tick virus that exhibited 100% identity with a virus from a dog rectal swab. Bambi tobamovirus was found in two libraries from fallow deer anal and mouth swabs, both libraries sampled from the same deer 2-4 weeks apart in a single NSW location and these sequences exhibited 99.8% nucleotide identity.

### Potential source and spread of tobamoviruses in Australia

As there are limited published sequences for RMV, TMGMV, and related viruses, it is challenging to determine when these viruses arrived in Australia and where they came from. The closest relatives to the Australian RMVs were sampled from Germany and Hungary (according to the CP phylogeny which has the most available sequence data, Figure 1B), while the closest relatives of the SG3-like tobamovirus viruses were from Hungary, Germany and New Zealand. However, these countries are not necessarily the source of the introduction but may have simply contributed a large proportion of the published tobamovirus sequences (particularly in the case of multiple RMV sequence from Germany). The Australian TMGMV clustered with viruses from Italy and Japan, based on the CP gene tree (Supplemental Figure 1), but based on the full genome phylogeny was most closely related to viruses from Germany. Accession OP525296, from Germany shared the highest identity with the virus detected here at 96.7% nucleotide identity across the complete genome, although there are a range of viruses from various countries that share more than 96% identity, including one from China (MZ395975), and Kenya (ON013908), that share 96.6% and 96.5% identity, respectively. Indeed, the most divergent TMGMV (MH730970) still shares 94.5% identity with the new Australian isolate. Considering the high levels of identity between isolates from different countries, it is not possible to get an indication of the source country. However, for each viral species, the Australian sequences clustered together (Figure 1A), suggesting that there was a single introduction of each of the five tobamoviruses into Australia, and subsequent spread to other states, although this is again subject to sampling biases.

### Abundance of plant transcripts in gut-metatranscriptomes

To assess potential hosts of the tobamoviruses detected in these libraries, we examined the metatranscriptomic data to identify transcripts from plants (Figure 2). The Poaceae family (grasses) were detected almost universally in the dog libraries isolated in QLD. This was the only plant family detected in three out of the five libraries sequenced and were only absent in one library, which was made up entirely of Fabaceae (legumes). Thus, the Poaceae and Fabaceae are strong candidates as hosts for bluey tobamovirus. Notably, grasses were present in most metatranscriptomes sampled in this study (Figure 2), albeit in lower proportions than in the dog libraries. The libraries from TAS, ACT and NSW contained a greater variety of plant transcripts making it harder to identify potential hosts (Figure 2). Both libraries containing bambi tobamovirus had transcripts from the Fabaceae, Plantaginaceae, and Poaceae, but among a range of many other potential hosts. Most RMV- and SG3-like tobamovirus-containing libraries prepared from samples sourced from herbivores (with the exception of Gudg-CC and 12-Kang2B) contained Plantaginaceae and/or Brassicaceae transcripts (among a range of other plant transcripts), which are known hosts for SG3 tobamoviruses (27, 29). However, the Plantaginaceae and Brassicaceae families were absent in the RMV-containing carnivore libraries, suggesting potential alternate hosts (Figure 2). There were no detectable plant transcripts in the tick library.

**Figure 2.**
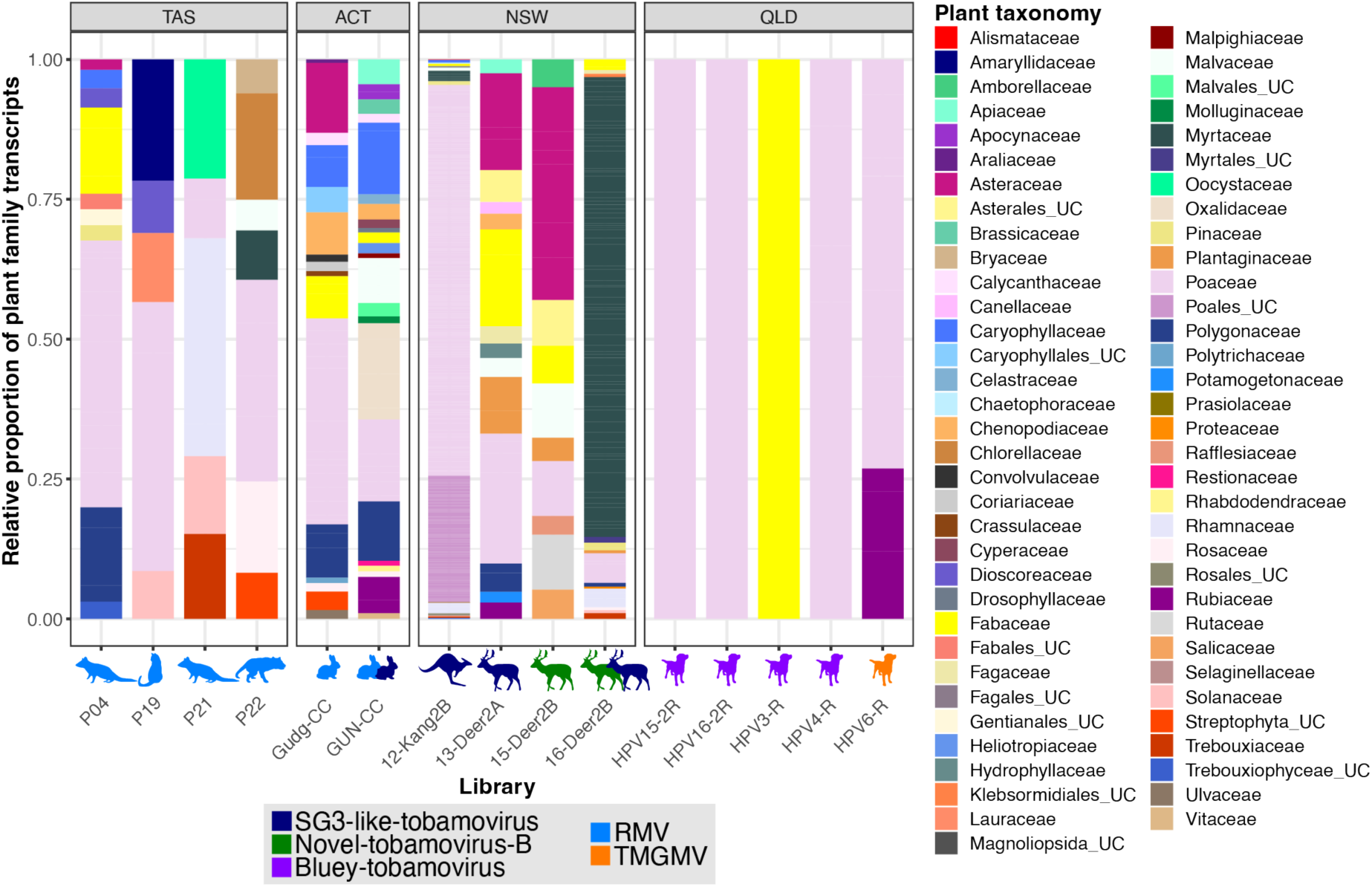
Abundance of plant transcripts in animal gut metatranscriptomes. Abundance of plant families is represented as a proportion of total plant transcripts present in each library. Animal silhouettes underneath each bar indicate the animal metatranscriptomes from which the plant transcripts were obtained and are coloured by viral species (purple=bluey tobamovirus; dark blue=SG3-like tobamovirus; blue=RMV; green=bambi tobamovirus; orange=TMGMV). Multiple animal silhouettes represent multiple viruses in that library. Note that libraries not pictured in the plot did not contain any plant transcripts.

### PCR confirmation

We employed RT-PCR of original RNA samples to verify the true presence of novel and exotic viruses in each individual library. The following viruses were detected in multiple libraries sequenced on the same lane: bluey tobamovirus, bambi tobamovirus, SG3-like tobamovirus, and RMV. RNA was not available for the ACT libraries, but for the libraries from NSW, QLD and TAS, the presence of the relevant viruses in individual RNA samples was determined by RT-PCR (except for libraries P19 and P21 for which there was insufficient RNA for all samples) (Table 3). Importantly, where RNA was available for testing, we confirmed the presence of all viruses detected in metatranscriptomes in one or more samples pooled in the library, including confirming the presence of the bluey tobamovirus in tick samples. While insufficient RNA precluded the use of RT-PCR to confirm that the RMV contigs in P19 and P21 are not the result of index hopping from other libraries, we determined that the contigs from these libraries are not identical to those from other libraries sequenced on the same lane, suggesting that RMV is indeed present in P19 and P21. Specifically, the P21 library had multiple RMV contigs which shared 96.4-99.4% nucleotide identity with contigs from P22, and 95.4-98.4% nucleotide identity with the single RMV contig from P04. P19 RMV contigs shared 97.6-98.4% and 97.8-98.1% nucleotide identity with contigs from P22 and P04, respectively. Equally, while RMV was detected in both of the libraries from the ACT that were sequenced on the same lane, the RMV contigs in these two libraries (Gudg-CC and GUN-CC) were not identical, sharing 98.4% nucleotide identity, and therefore RMV is likely present in both libraries.

**Table 3.**
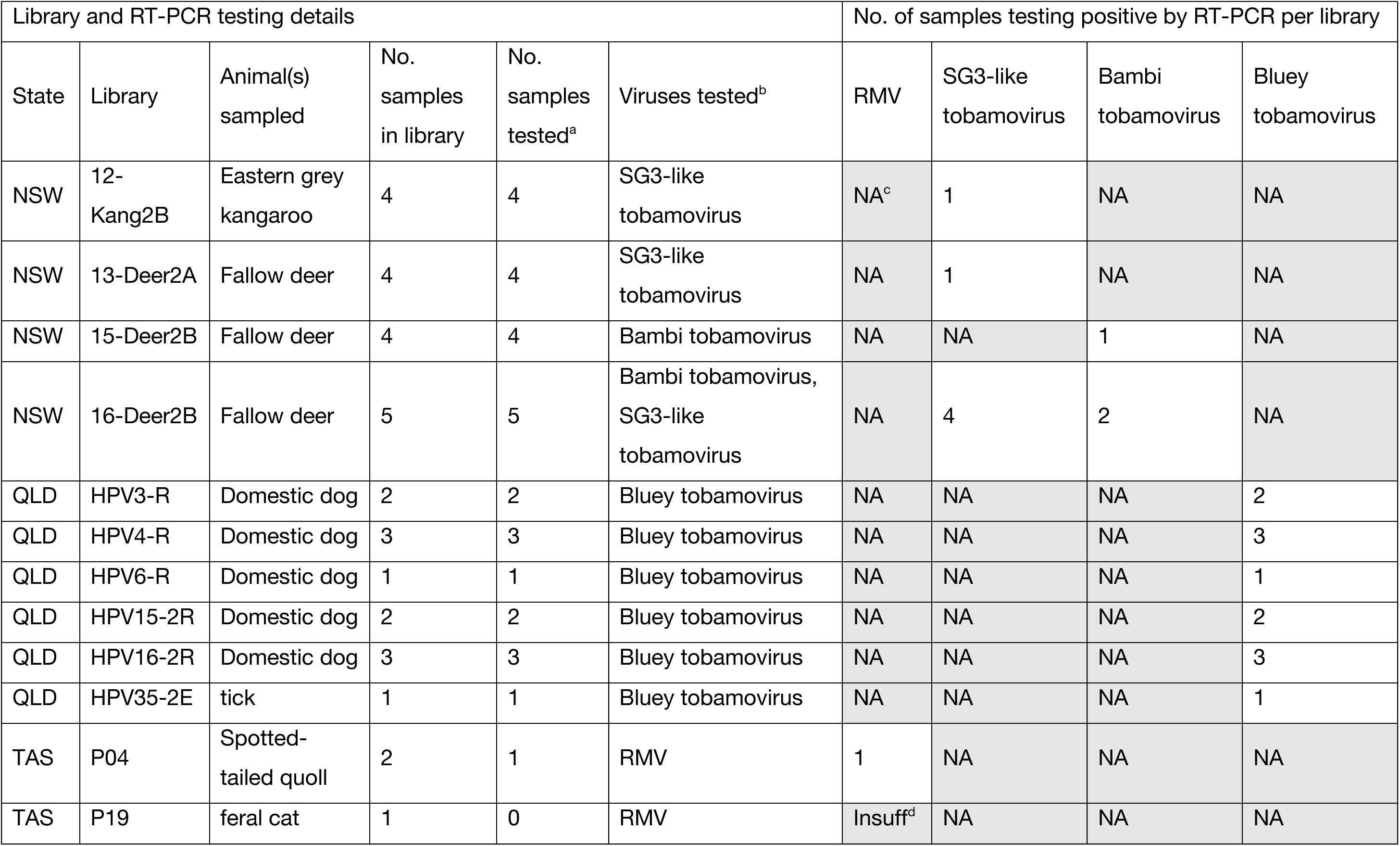
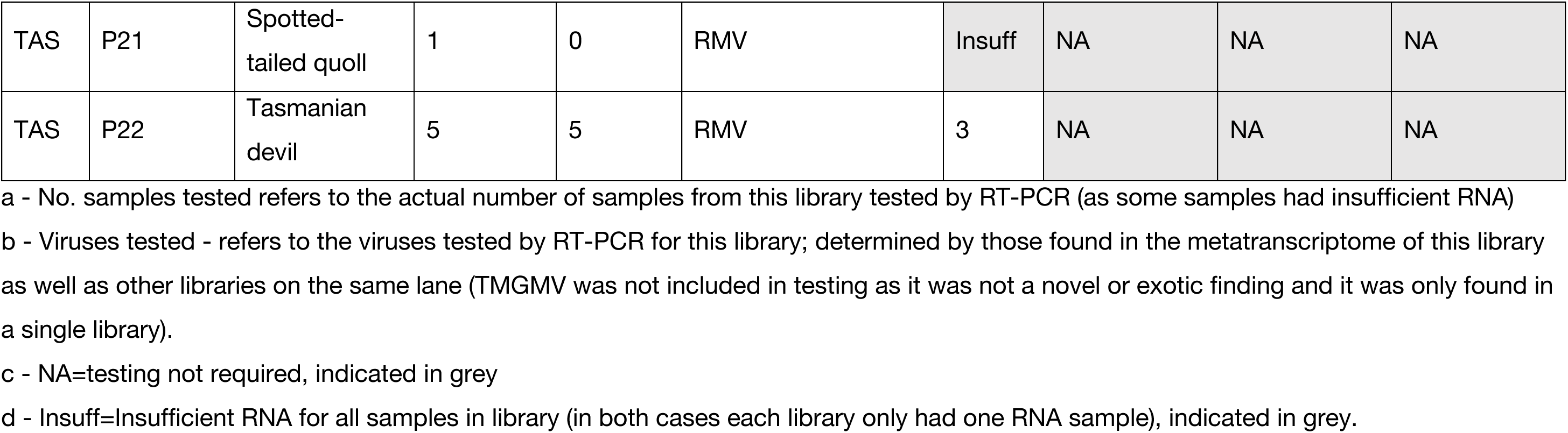
RT-PCR confirmation of detected viruses.

## Discussion

Tobamoviruses are known to have devastating impacts on agricultural and horticultural crops. Consequently, they are listed as biosecurity threats in many countries, including Australia, where many tobamoviruses are not known to be present. Australia imports seeds for the horticultural industry under stringent phytosanitary inspections. However, with increasing climate change and globalisation, Australia remains vulnerable to the introduction of seed or vector transmitted viruses (46, 47). Surveillance has been used in Australia to monitor virulent viruses and their vectors and has been critical for the early detection of exotic viruses, as well as demonstration of freedom from exotic or endemic pathogens (47). The surveillance methods vary by region, but mainly involve the use of targeted molecular approaches and immunoassays applied to survey particular field crops and samples at the border (32). These approaches are limited in that they are unlikely to detect unknown viruses and potential endemic variants, and may fail to detect incursions in a timely manner (15, 48). In addition, viruses may invade or persist in wild or native host vegetation habitat before spilling over to cultivated land. As such, combining surveillance in field crops and border samples with metatranscriptomic surveillance of wild and domestic animal gut material, offers an alternative virus surveillance strategy for early detection of exotic virus incursions, and demonstration of eradication or containment measures. In this study, metatranscriptomic sequencing of opportunistically sampled wild and domestic animal gut material in Australia revealed the presence of five different tobamovirus species, including one that was thought to be exotic (RMV) and three novel viruses (a *tentative* novel relative of subgroup 3 tobamoviruses – SG3-like tobamovirus – and two highly diverse novel tobamoviruses, bluey tobamovirus and bambi tobamovirus). This is the first study to explore the use of metatranscriptomic sequencing of animals as a strategy to detect exotic crop viruses.

The significance and potential impact of the presence of each of the five tobamoviruses detected here varies. The detection of RMV in Australia in native and feral animal gut metatranscriptomes is critical as this virus was previously considered to be exotic in Australia. Likewise, the detection of the SG3-like tobamovirus is also important, as its closest relatives, RMV and TVCV are known plant pathogens (27). Furthermore, this virus clusters within the larger subgroup 3 clade of viruses containing RMV, TVCV, WMoV, and YoMV, which are all important pathogens of Brassicaceae and Plantaginaceae (27, 49–52). Therefore, it is highly possible that this virus could be pathogenic in Brassicaceae and Plantaginaceae plants like its subgroup 3 relatives (27, 49–52) as transcripts from these host families were found in three of four libraries containing SG3-like tobamovirus virus. In addition, the New Zealand virus that clusters with the SG3-like tobamovirus based on CP sequence (HQ389333; originally characterised as TVCV), was found in a *Plantago* species presenting with mild chlorosis or mottles (27). Nevertheless, a broader survey of the SG3-like tobamovirus species in various hosts should be conducted in the future, along with virus host studies.

The detection of TMGMV is not alarming as this virus is among a number of tobamoviruses that have previously been detected in Australia (53), including CGMMV (14), yellow tailflower mild mottle virus (YTMMV) (54), clitoria yellow mottle virus (55), YoMV (48), pepper mild mottle virus (48), hibiscus virus (AY664875), tomato mosaic virus (NC_002692), odontoglossum ringspot virus (KF855954.1), and tomato mottle mosaic virus (56). Indeed, TMGMV has been present in Australia since the late 1800s (53). TMGMV seems well adapted to members of *Nicotiana* (57) and is particularly common in wild *Nicotiana glauca* wherever it is present around the world, including Australia (53). However, TMGMV is known to infect a wide range of hosts, including members of the Solanaceae, Umbelliferae, Gesneriaceae, Rubiaceae and Poaceae families (28, 58), and transcripts of the latter two were present in the TMGMV containing library, suggesting these as potential hosts in this case.

It is difficult to speculate on the pathogenic potential or likely hosts of the two highly novel viruses detected in this study – bluey tobamovirus and bambi tobamovirus – because they are both distinct from known viruses. However, analysis of the eukaryotic transcripts in the libraries containing bluey tobamovirus and bambi tobamovirus viruses suggests grasses (Poaceae) and potentially legumes (Fabaceae) as strong candidate hosts for bluey tobamovirus and bambi tobamovirus, along with Plantaginaceae for bambi tobamovirus. It is also possible that the true natural host for these viruses were not detectable in the metatranscriptome due to low abundance or RNA degradation, as eukaryotic RNA transcripts would not be protected like encapsulated viral RNA. Indeed, there were no detectable plant transcripts in the tick library (which contained bluey tobamovirus).

As these viruses have not been detected elsewhere, is it possible that the natural hosts of bluey tobamovirus and bambi tobamovirus are wild Australian native plants, as previously shown for yellow tailflower mild mottle virus (54). As a tobamovirus originating from native Australian plants has been shown to naturally spread to exotic plant species (22), a broader understanding of the distribution and the natural hosts of these viruses could be beneficial for both native conservation, and their management to deter potential spillover or spread to cultivated crops. Notably, there was no evidence that any of the tobamoviruses detected in animal metatranscriptomes are present in horticultural farms in Australia. Thus, increased field surveillance efforts remain vital to protect the horticultural industry.

We detected these viruses in a variety of animal guts, including native and feral wild-living species and domestic animals, and assume that the viral hosts are plants within the animal diet. Interestingly, we also found some of these viruses in carnivores and hypothesize that the viruses were in plants that were incidentally consumed or in the gut of their ingested prey. This is possible due to the high stability of tobamoviruses and their ability to remain viable in the environment for a long time (30, 31). As tobamoviruses are spread by mechanical transmission (30), it is likely that vertebrate species may play a role in the spread of these viruses. It has already been shown that plant viruses may colonize new environments through grazing animal teeth, ingestion of plants, and subsequent passing of the viruses through the gut (24). As such, these data emphasize the potential ecological importance of wild animals in the spread of tobamoviruses and other contact transmitted viruses.

Interestingly, bluey tobamovirus was found in high abundance in a tick metatranscriptome with a near identical viral genome sequence (99.8-100% nucleotide identity) to those found in dog faeces. As this virus is highly divergent from known tobamoviruses and no plant transcripts were found in the tick library, we cannot rule out an animal tropism. However, since no tobamoviruses are known to be arboviruses, it is unlikely that this virus was replicating in ticks or dogs. Further, it is not uncommon for plant viruses to be detected in (and transmitted by) invertebrates (24, 26, 48). Indeed, 80% of plant viruses rely on insect vectors for mechanical transmission (59), and invertebrate pollinators (such as the honeybee and bumblebee) have been shown to transmit tobamoviruses (60, 61). As ticks are not herbivores, it is unlikely that they would play a major role in plant virus transmission. However, ticks are commonly found in plants, as they climb into vegetation to be in a position to fall on passing animals. It is therefore possible that the bluey tobamovirus mechanically adhered to tick body parts following tick contact with infected sap, through physical abrasion of an infected plant, as known to occur with plant viruses transmitted by bees and other pollinator insects (60–63). As such, ticks could play a role in wider dispersal of tobamoviruses. Indeed, a number of tobamoviruses were found in tick viromes in China, including Tobacco mosaic virus, YoMV, and CGMMV (64). As the dog and tick sampling was done at an outdoor pop-up vet clinic in regional Queensland, it is also possible that the dog and tick samples were contaminated at the point of sampling. Regardless of the host origins or transmission pathways of these viruses, ectoparasites should also be considered during the management and control of these viruses in the field.

These findings demonstrate that total RNA-sequencing of wild and domestic animal gut swabs and faecal material is a potentially useful alternative strategy for the surveillance of economically important plant viruses in Australia, in addition to targeting specific field crops or natural plant hosts. This work also demonstrates that tobamoviruses are detectable in a range of different animals (including domestic, native and introduced wild animals, herbivores, carnivores, and even invertebrates) and that any method of sampling the gut content (i.e. anal/rectal swab, faecal sample, gut content sample) can be applied. While current surveillance methods involve direct sampling of individual plants or seed lots, gut metatranscriptomics allows a larger scale sampling approach whereby a pool of animal faeces, representing an entire range of ingested plants can be sequenced. This may reduce the labour involved in the field sampling effort. In addition, where RNA extraction from plant material can be challenging, the plant material in animal gut has already been broken down, and therefore extraction from animal material could be a simpler process requiring less complex tissue disruption methods. Conversely, a limitation of the gut metatranscriptomic approach is that viruses that are not highly stable in the environment may be less likely to be detected (24) due to degradation within the animal gut. It may also often fail to detect DNA viruses as only RNA transcripts from these viruses are expected to be sequenced and these may be degraded in the animal gut since they would not be protected by capsids. However, considering RNA viruses are the most abundant plant pathogens, causing nearly one-third of global crop losses (65), this approach could be a highly beneficial addition to current surveillance techniques, enabling detection of a broad range of potential viral threats. In particular, animal gut metatranscriptome-based surveillance would be a valuable tool for detection of potential viral threats circulating in the wild, facilitating mitigation techniques to avoid spill over into cultivated crops. Interestingly, in Australia, metatranscriptomics has also been employed to detect plant viruses on honeybees (48), including the devastating CGMMV known to have caused major losses in the horticultural industry (66). Although CGMMV was first detected in Australia from a survey of cucurbit crops in the Northern Territory conducted in July 2014 (14), metatranscriptomic surveillance of honeybees detected low levels of CGMMV in additional states, Queensland and Western Australia, prior to its identification in field crops (48). This further demonstrates the utility of metatranscriptomics of animals for plant virus surveillance.

Due to a lack of global sequence data for RMV and related viruses, it is difficult to determine their source of introduction into Australia. The TMGMV detected in this study is most closely related to viruses detected in other countries, including Germany, Japan and Italy. However, this does not necessarily indicate that the detected virus was recently introduced from overseas as TMGMV has been endemic in Australia since the early 1900s (53) and is known to demonstrate little genetic variation over time (53). However, our work indicates that there has been at least one recent incursion event of a known virus – RMV. As the Australian RMV sequences formed a monophyletic group, the current evidence suggests that there was a single incursion of this virus. The SG3-like tobamovirus also seems likely to be from a single introduction and, based on CP sequence, is closely related to New Zealand and German sequences (although support for the German and New Zealand viruses’ position in the CP tree was weak). Finally, the two highly diverse novel tobamoviruses, bluey tobamovirus and bambi tobamovirus, are so distinct from known viruses that it is difficult to determine if they represent exotic incursions or whether they are endemic in Australian plants, like YTMMV (54).

Importantly, tobamoviruses were detected in animals in four Australian states and territories, and RMV and SG3-like tobamovirus were each found in two states and territories indicating spread of these viruses within Australia. RMV was detected in both the ACT and Tasmania, even though these locations are more than 700 kms apart and separated from the mainland of Australia by Bass strait, >200km of water. Viruses may have spread over Bass strait through their ability to be contact transmitted via fomites (15) – potentially on shoes or equipment travelling by air or on car tyres on ferries, or even by migratory birds. Future studies should focus on understanding the distribution and the impact of RMV and the other tobamoviruses detected here.

In sum, we determined that metatranscriptomic screening of animal gut content could serve as a non-targeted surveillance tool, particularly in conjunction with existing methods of plant biosecurity surveillance. Future work should focus on characterizing the host range, geographic distribution and the impact potential of the new and known tobamoviruses detected here. The significant distribution of these viruses in Australia demonstrates the importance of early detection to prevent cryptic spread. This study also solidifies the importance of continuous integration of innovative rapid diagnostic and surveillance genomics tools to support viral disease management for sustainable food production.

## Acknowledgements

We thank Wendy Forbes assisting with samples submission to AGRF and ordering the positive controls. We acknowledge the University of Sydney’s high-performance computing cluster Artemis for providing the computing resources used for this study.

