## Supplementary Table 1 and Supplementary Figure 1 for "Detection of exotic biosecurity threat ribgrass mosaic virus and novel tobamoviruses through metatranscriptomic sequencing of animal gut content"

**Table S1. Primer sets used for RT-PCR confirmation of viruses and amplification and Sanger sequencing of SG3-like tobamovirus**

| Primer name | Primer sequence 5'-3' | Direction | Target | Purpose | Amplicon size |
|---|---|---|---|---|---|
| **Full genome amplification and Sanger sequencing of SG3-like tobamovirus** | | | | | |
| FP1b | ACAAACAACAACAACATGGC | Forward | SG3-like tobamovirus | RT-PCR/Sanger | 2,003 bp |
| RP1 | AACTTGTCGAAACCACCACCT | Reverse | | RT-PCR | |
| FP2 | CGACGGAGGAGGAGATTTCAG | Forward | SG3-like tobamovirus | RT-PCR/Sanger | 1,484 bp |
| RP2 | GACCCCTGCTTCGACTTTGT | Reverse | | RT-PCR | |
| FP3 | GACGTACGAGAAGACTGCGA | Forward | SG3-like tobamovirus | RT-PCR/Sanger | 2,106 bp |
| RP3 | CGGTTTCCTCTTAGCGTCCA | Reverse | | RT-PCR | |
| FP4 | ACTTCTGCGGTCGTTACGTT | Forward | SG3-like tobamovirus | RT-PCR/Sanger | 1,723 bp |
| RP4 | CGGGGTTAGGGAGGATTCGA | Reverse | | RT-PCR | |
| FT1 | AGCGCTACTCAGAAAGAACGT | Forward | SG3-like tobamovirus | Sanger | N/A |
| FT2 | AAGTTGGGTCATGTGCAGGA | Forward | SG3-like tobamovirus | Sanger | N/A |
| FT3 | ACATGGTTTAGGGTGGCTGT | Forward | SG3-like tobamovirus | Sanger | N/A |
| FT4 | ACAGTTGGTCAGTTAGCGGA | Forward | SG3-like tobamovirus | Sanger | N/A |
| FT5 | TGGACGCTAAGAGGAAACCG | Forward | SG3-like tobamovirus | Sanger | N/A |
| FT6 | AGGGTCCCGGGTGTATGTTA | Forward | SG3-like tobamovirus | Sanger | N/A |
| **Detection RT-PCR** | | | | | |
| RMV_410_F | GCGATAAGTGATCCGGACGT | Forward | RMV | RT-PCR | 565 bp |
| RMV_974_R | ACTTCTAATGGCGACGGTCG | Reverse | | | |
| F_Bamb | AGATTGGAGAAGGGTTGCGG | Forward | Bambi tobamovirus | RT-PCR | 540 bp |
| R_Bamb | TCAGCAGGACATCGCAAAGT | Reverse | | | |
| F_Blue | CTACGTGTAGCCGTCTCGAC | Forward | Bluey tobamovirus | RT-PCR | 226 bp |
| R_Blue | TGCGAATCATTTCAGCAGCG | Reverse | | | |
| F_Novel-SG3 | AATTGGAGGAAGGGATGTGGTT | Forward | SG3-like tobamovirus | RT-PCR | 571 bp |
| R_Novel-SG3 | AAGCCTCAAACTCTGCCYTG | Reverse | | | |

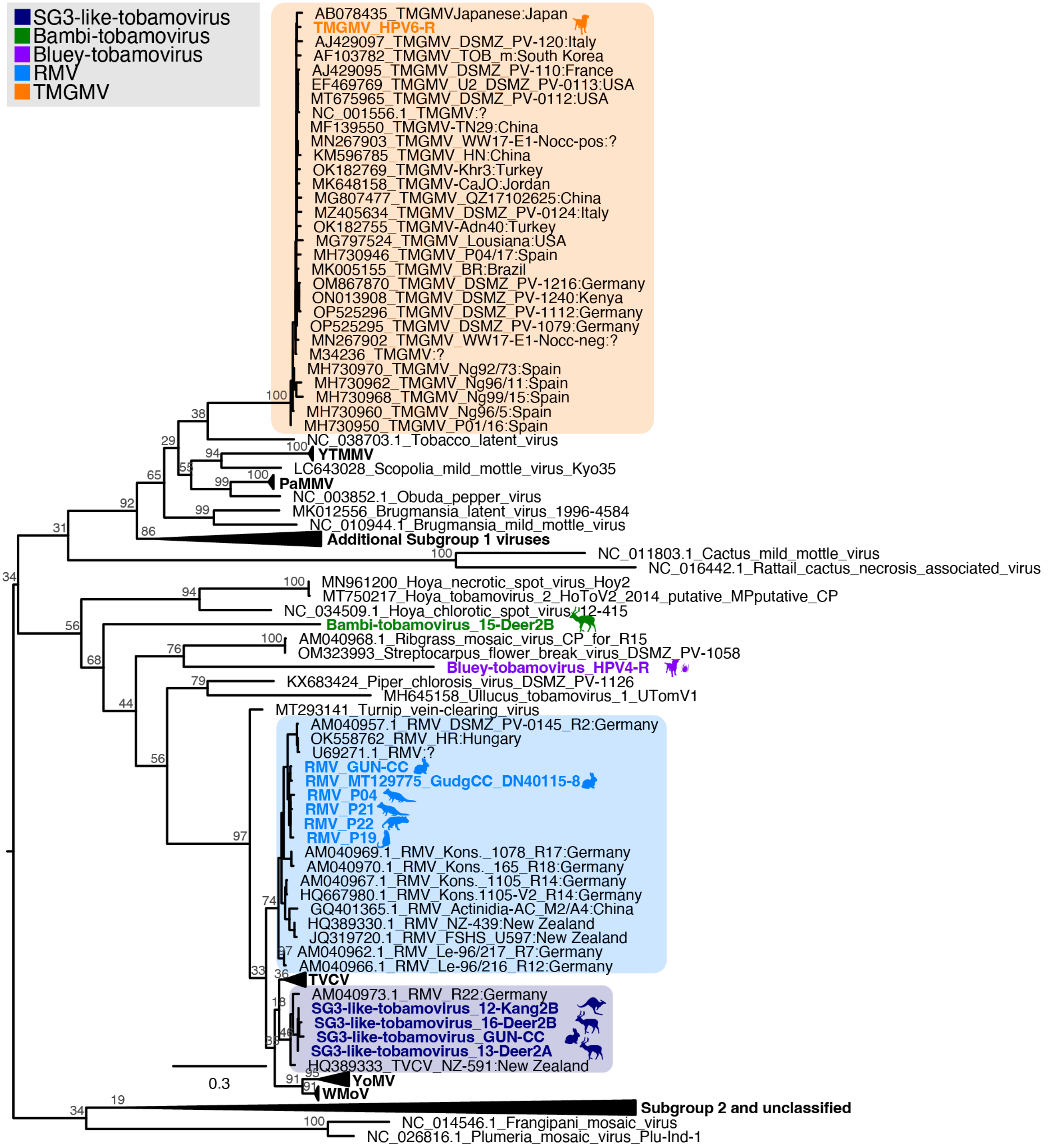


**Figure S1. ML phylogenetic tree of the coat gene sequences of tobamoviruses detected in Australian animal metatranscriptomes together with published coat sequences.** Sequences obtained from animal metatranscriptomes in Australia are indicated in bold and coloured by viral species, and the clades that they cluster within are highlighted in the same colour: purple=bluey tobamovirus; dark blue=SG3-like tobamovirus; blue=RMV; green=bambi tobamovirus; orange=TMGMV. Animal silhouettes beside the clades indicate the animal metatranscriptomes from viruses of that species were obtained and are coloured by viral species. For viruses with 100% identity in the coat gene, only a single taxon was included as a representative, but multiple animal silhouettes may be used to indicate the range of metatranscriptomic sources. The GenBank accession number for published sequences is indicated at the start of the taxon name. Numbers at the nodes indicate the percentage support from 1,000 bootstrap replicates and the trees are midpoint rooted. The location (country) of collection is indicated in the taxa name in relevant clades.
